# Automated Microfossil Identification and Segmentation Using a Deep Learning Approach

**DOI:** 10.1101/661694

**Authors:** L.E Carvalho, G. Fauth, S. Baecker Fauth, G. Krahl, A. C. Moreira, C.P. Fernandes, A von Wangenheim

**Affiliations:** Graduate Program in Computer Science - Federal University of Santa Catarina - Campus João David Ferreira Lima - Trindade - Department of Informatics and Statistics - Room 320 Florianópolis/SC – CEP 88040-97; Image Processing and Computer Graphics Lab - National Brazilian Institute for Digital Convergence.; itt Fossil – Instituto Tecnológico de Micropaleontologia, Universidade do Vale do Rio dos Sinos, Av. UNISINOS, 950, 93022-000 São Leopoldo, RS, Brazil; Graduate Program in Materials Science and Engineering Federal University of Santa Catarina, Florianópolis, SC, Brazil

## Abstract

The applicability of computational analysis to paleontological images ranges from the study of the animals, plants and evolution of microorganisms to the simulation of the habitat of living beings of a given epoch. It also can be applied in several niches, such as oil exploration, where there are several factors to be analyzed in order to minimize the expenses related to the oil extraction process. One factor is the characterization of the environment to be explored. This analysis can occur in several ways: use of probes, extraction of samples for petrophysical components evaluation, the correlation with logs of other drilling wells and so on. In the samples extraction part the Computed Tomography (CT) is of importance because it preserves the sample and makes it available for several analyzes. Based on 3D images generated by CT, several analyzes and simulations can be performed and processes, currently performed manually and exhaustively, can be automated. In this work we propose and validate a method for fully automated microfossil identification and extraction. A pipeline is proposed that begins in the scanning process and ends in an identification process. For the identification a Deep Learning approach was developed, which resulted in a high rate of correct microfossil identification (98% of Intersection Over Union). The validation was performed both through an automated quantitative analysis based upon ground truths generated by specialists in the micropaleontology field and visual inspection by these specialists. We also present the first fully annotated MicroCT-acquired publicly available microfossils dataset.

## Introduction

The applicability of computational image analysis to paleontological data encompasses the possibility of identifying, reconstructing and visualizing microfossils in rock samples not recovered by traditional extraction methodologies. It can also allow the taxonomic microfossil identification even before the physical extraction from the rock sample. In addition, it is also pertinent to verify the microfossil position in a given sedimentary stratum, which can help in taxonomic inference, whereas detailed positional information is lost in the traditional preparation method [8]. Computational analysis of samples can be applied in several niches, e.g. oil exploration, habitat reconstruction and geology and paleontology research.

On the other side, in the oil exploration field, there are many factors to be taken into consideration in order to minimize oil prospecting expenses. One factor are the environmental conditions, which can be analyzed in multiple ways: use of probes, extraction of samples for petrophysical components evaluation and correlation with logs from other drilling wells.

In the area of samples extraction it is possible to perform different analyzes on a given sample. Here Computed Tomography (CT) plays a central role. More specifically, samples are analyzed with X-ray micro-tomography (MicroCT), which is a radiographic imaging technique that produces 3D images of the material’s internal structure with a spatial resolution of around 1 micrometer [11]. MicroCT is of significance because it preserves the sample and makes it available for different studies. Based on MicroCT generated data volumes, various 3D data analyzes and simulations can be performed and several analysis processes can be computationally carried out and automated using state-of-the-art Computer Vision (CV) techniques. These processes are currently performed manually and in a time-consuming manner. One of these processes that can undergo automation through CV is the microfossils identification and localisation in rock samples, which is the focus of this study.

### Objective and Strategy

In this work, we propose a CV workflow composed of computational methods that starts with the MicroCT scanning process of a sample and ends with the fully automated identification and localisation of individual microfossils in this sample. The main research question we try to answer is: *Is it possible to fully automatically and reliably identify microfossils in carbonatic rock samples?*

The novelty in our approach is the use of Deep Learning Convolutional Neural Network (CNN) approaches for the identification and 3D segmentation of microfossils directly in their deposition place. Our approach works directly on MicroCT data gained from carbonatic rocks, without the need of any preparation or physical extraction. For this purpose we developed an identification and segmentation strategy that employs a special category of CNN models, namely Semantic Segmentation (SS) neural networks and extends this model in order to be able to process whole 3D MicroCT sample volumes. In order to identify the best model, we extend, train, test and compare a series of different state-of-the art SS models. For the validation our approach we employ a validation strategy where we compare our results to ground truths that were manually generated by experienced micropaleontologists employing state-of-the-art automated image segmentation validation algorithms.

### State of the Art

Paleontology is a well-established science and its methodological intersection with the computational field started to grow in the 1990’s. In the late 1980’s, most main paleontology journals still show an irregular presence of computational methods: some journal issues contained one article describing some computational method application, others presented 2 or 3 articles and very few offered a larger number of them [24]. In the majority of journals and books the insertion of computational methods in the paleontology field still looked uneven.

In the late 1990s, however, with the widespread use of medical CT, a growth in research activities employing tomographic images occurred [24]. This boosted the development of specialized software applications such as: DRISHTI, SPIERS, SEG3D, IMAGEJ, MIMICS, VGSTUDIO MAX, AVIZO, AMIRA, Geomagic, Rhinoceros, Imaris, ITK-SNAP and TurtleSEG. These specialized tools helped change how researchers deal with specific problems in several fields, including geology and paleontology, frequently with applications to oil and gas exploration. The applicability of the set of tools and techniques that came to be called Virtual Paleontology (VP) range from animal, plant and microorganisms evolution analysis until the virtual reconstruction of a specific extinct environment [23].

On the other side, the application of the study of microfossils to the area of oil prospection had its first appearance in 1890 in Poland [21], but it was in the USA, in 1920, with the use of microfossils to identify the age of probes extracted from drilling rigs, that a bigger advance in the development of the field of Applied Micropaleontology was attained [14].

In the last decade multiple research works contributed to improve the micropaleontology field. The latest efforts aim at the use of VP associated with CNNs in order to identify microfossils [5]. With this in mind, our research is focusing on pursuing techniques that can identify microfossils on their deposition place, *i.e.*, without the need of previous physical isolation. For this purpose we research some CV fields such as 3D segmentation applied to tomographic image and 3D object recognition, in order to apply them to microfossil identification.

In the next subsections we summarize the results of the systematic literature reviews (SLR) we performed in order to identify the state-of-the-art of the methods and procedures that potentially could be used in microfossil image studies. These reviews followed the approach originally proposed by [10] for SLRs in Computer Sciences, where first we defined a research question: *Is it possible to fully automatically and reliably identify microfossils in carbonatic rock samples?*. This broad question, in order to be more manageable, was split into 2 topics, each of which was explored in depth in a separate SLR:

- Analysis of 3D segmentation methods applied to tomographic images, which could possibly be used to segment microfossils [3];
- Analysis of methods used for 3D object recognition in a general context, aiming to evaluate which methods could be applied to the microfossils field [4].

The results of these two SLRs will be briefly summarized below. Since a detailed description would exceed the scope of this paper, we refer to the referenced SLRs for more details.

### 3D segmentation applied to tomographic images and 3D object recognition

An initial analysis of image processing methods employed in the fossil identification area showed difficulty in finding any works that explore microfossils. So we generalized our search for methods in other similar areas. We started by performing a systematic literature review on 3D segmentation methods applied to tomographic images [3]. Several works were analyzed which comprehended a vast group of segmentation methods. In our review, we noticed a tendency on the use of 3D segmentation methods based on models and region growing. However, its use for fossil/microfossil segmentation wasn’t noticed in the literature.

We also analysed the field of 3D object recognition employing the same SLR methodology [4]. In this SLR for 3D object recognition we could identify two general pipelines. Both pipelines start with the data acquisition, which can basically vary between 3D data (MRI, CT) or 2D data (RGB and RGBD cameras), pre-processing, where methods for artifact removal, image enhancement and image simplification and data representation, wherein several authors proposed a varying amount of different object representations. Then, it comes the stage where both pipelines differ: In the first pipeline, the data representation stage is used to describe and storage the object representation chosen, which is later used for similarity calculation and object identification; In the second pipeline, the data representation is employed for training a specific recognition architecture, such as a CNN, which is afterwards used for other objects recognition. Despite having found two general approaches for 3D object recognition, we could not identify, in our review, the application of these approaches on fossil identification.

### Deep learning, object recognition and paleontology

The 3D object recognition area has, in the last few years, experienced a growing boosted by the increased availability of new algorithms and models, 3D data and the popularization of a varied palette of 3D sensors. Methods developed in this area find application in a wide range of areas, from the field of robotics to the security and surveillance domain. The general tendency in this area has been the use of Deep Learning (DL) techniques.

DL is a form of machine learning that enables computers to learn from experience and understand the world in terms of a hierarchy of concepts [6]. DL employs very deep CNNs, with neural networks that sometimes consist of more than 100 layers, in contrast to the Artificial Neural Networks (ANNs) employed between the 1980’s and 2000’s, that typically employed only three layers. One key concept here is the Convolutional Layer (CL), a feature extraction structure, first presented in [12], that allows the hierarchical learning and representation of complex knowledge. Because DL CNNs gather knowledge from examples, there is no need for a human computer operator to formally specify all the knowledge that the computer needs. The capacity to represent a hierarchy of concepts in a network dozens of CLs deep allows a DL CNN to learn complicated concepts by building them out of simpler ones; a graph of these hierarchies would be many layers deep [6].

One work that employs DL for object recognition is the 3D Object Recognition with Deep Belief Nets approach [15], where a network of symmetrically connected neuron-like units, that performs stochastic decisions about whether to be on or off, is presented. In the Convolutional-Recursive Deep Learning for 3D Object Classification, where Socher [22] presents a model based on the combination of convolutional and recursive neural networks for the feature learning and classification in RGB-D images. Another approach is the Vision-based Robotic Grasping System Using Deep Learning for 3D Object Recognition and Pose Estimation, where Jincheng Yu [30] presents a robotic vision-based system, which can not only recognize different objects, but also estimate their pose through the use of a deep learning model. The deep learning model used is the Max-pooling Convolutional Neural Network (MPCNN).The 3D Object Recognition and Pose estimation System Using Deep Learning Method is an approach where Dong Liang [13] presents a 3D object recognition and pose estimation method using a deep learning model. Recognizing multi-view objects with occlusions using a deep architecture, where a method for efficient 3D object recognition with occlusion is presented [26]. In the Convolutional neural network for 3D object recognition based on RGB-D dataset, Jianhua Wang [25] employs a convolutional neural network model to learn features from a RGBD dataset which are given to a linear SVM to classify objects. In the Convolutional Neural Network for 3D object recognition using volumetric representation Xiaofan Xu [29] presents an efficient 3D object volumetric representation, called Volumetric Accelerator (VOLA), which requires much less memory than a normal volumetric representation. Properly, VOLA can reduce the computational complexity of Convolutional Neural Networks (CNNs). None of these approaches tackles the problem of identification of fossils embedded in rocks or any remotely similar problem.

## Material and Methods

This section describes our datasets and the CV approach we developed for fully automated microfossil identification and segmentation in carbonatic rock samples.

### Material

We employed two datasets: a scanned carbonatic rock sample obtained from a drilling rig probe and a set of manually isolated microfossil specimens that were afterwards obtained from this sample. The sample was collected at the Sergipe Basin Quaternary sediments (Fig. 1):

- A *carbonatic rock sample* was the material for which we developed our CV approach. The MicroCT scanner used to digitise the sample is a Versa XRM-500 (ZEISS/XRadia) with the following specifications: best resolution (pixel size) 0.7 μm, voltage 30-160 kV, power 2-10 W, CCD cameras 2048×2048 pixel, optical lenses 0.4X, 4X, 10X, 20X and 40X, a set of 12 filters for beam hardening correction, maximum sample mass capacity 15 kg and sample size limit (diameter / height) 80*/*300 mm. The sample acquisition parameters we employed are: Spatial resolution 1.08 mm, image size 956×1004×983, no filtering for beam correction hardening, 10X optical lens, 30 kV / 2W, angular step 0.255 (1600 projections) and exposure time 11 seconds. Figure 2 shows the rock sample and an excerpt of one slice of its digitised result.
- A set of *manually isolated microfossil specimens*, gained from the sample above, was employed in this work for illustration purposes and as a guide in order to allow us to know how the specimens in the *rock sample* would look like if cleanly segmented. These microfossils were prepared in the laboratory, following specific precautions so that there were no chemical and/or mechanical changes: (i) the sediment was first immersed in deionized water for approximately 24hrs, aiming the chemical disaggregation; (ii) then, it was washed with running water in a 63 μm sieve; (iii) next, the material was dried at 40*degree*C for approximately 48 hours. After drying the samples, the main representative microfossils in the sample were selected through a magnifying glass. In this work, the microfossils specimens were stamped with the help of a multidimensional acquisition with the Zeiss Discovery V20 stereoscope (Z-stak mode in AxioVision 4.8 software). Figure 3 presents these microfossils.

**Figure 1.**
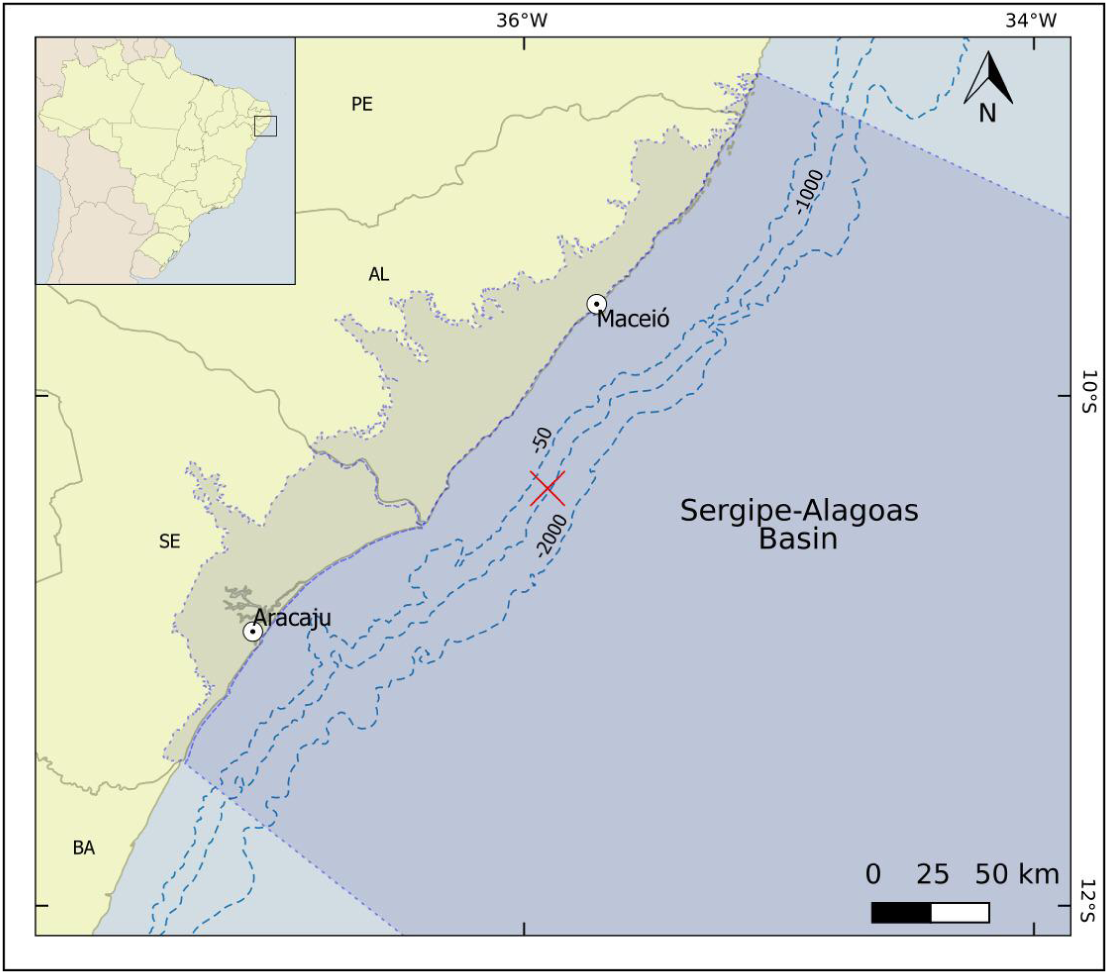
Sergipe Basin map with the exact rock samples extraction location marked with a red cross. The sample was collected at a depth of approximately 2,500 meters. Source: the authors.

**Figure 2.**
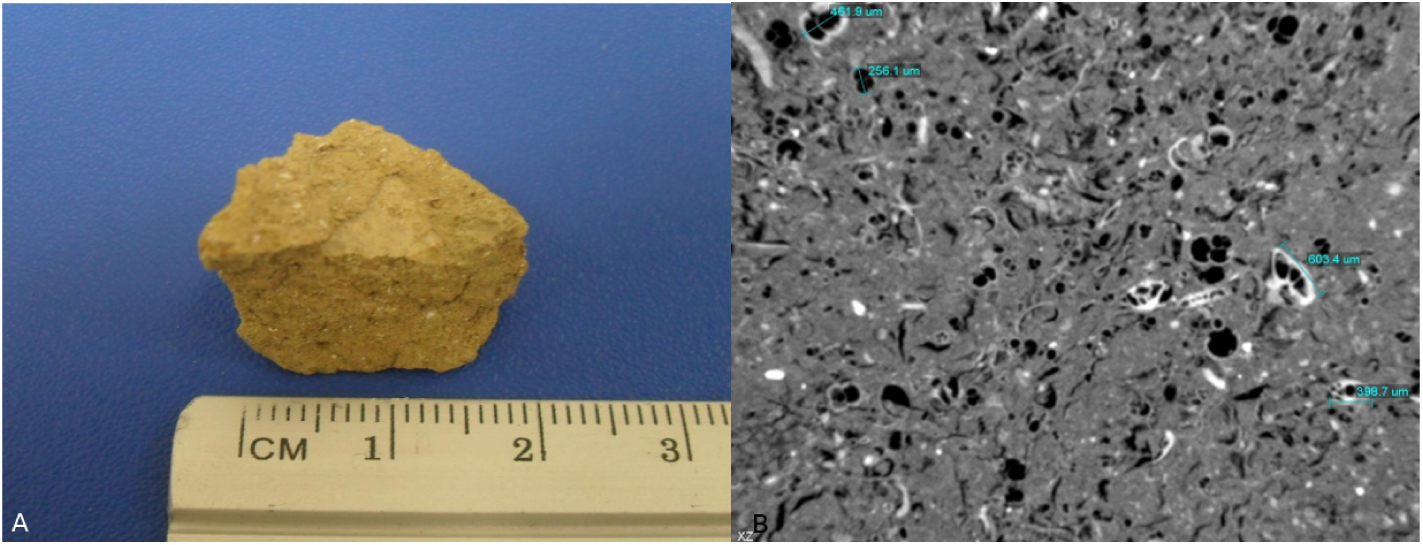
Analyzed rock sample (A) and one of its microtomography 2D sections image (B). Source: the authors.

**Figure 3.**
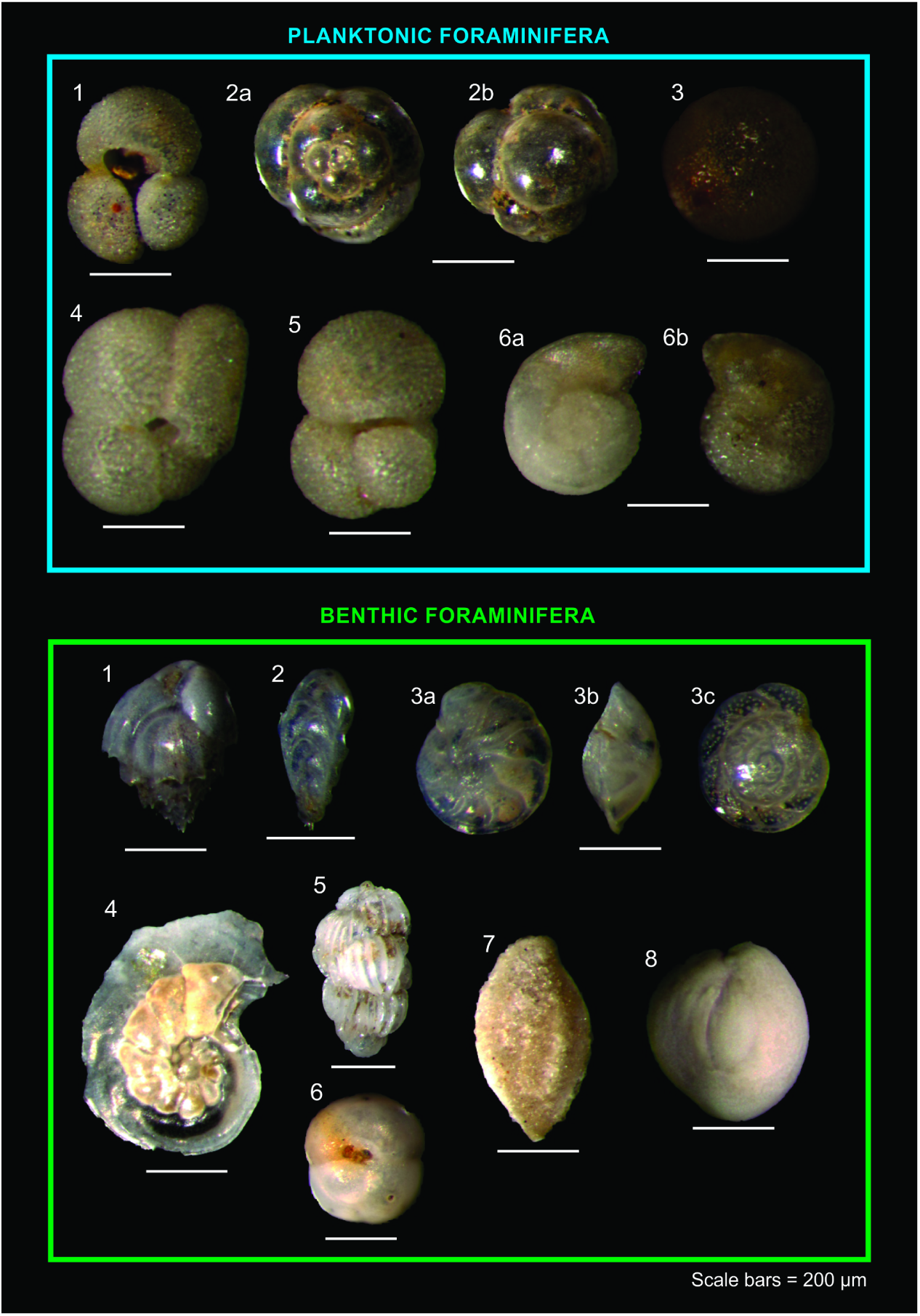
Analyzed foraminifera specimens. Planktonic Foraminifera:1) Globogeri-noides ruber; 2a-b) Candeina nitida; 3) Orbulina universa; 4) Globigerinoides trilobus saculifera; 5) Globigerinoides trilobus; 6a-b) Globorotalia truncatulinoides. Genus of benthic foraminifera: 1) Bulinina; 2a-c) Bolivinita; 3a-c) Cibicidoides; 4) Laticarinina; 5) Uvigerina; 6) Sphaeroidina; 7) Siphonaperta; 8) Quinqueloculina. Source: the authors.

The dataset containing the MicroCT data and and the manually segmented images annotated by specialists is available at: http://www.lapix.ufsc.br/microfossil-segmentation

### Methods

The CV approach we present here is intended to be embedded into a broader workflow. Figure 4 presents a general overview of this workflow.

**Figure 4.**
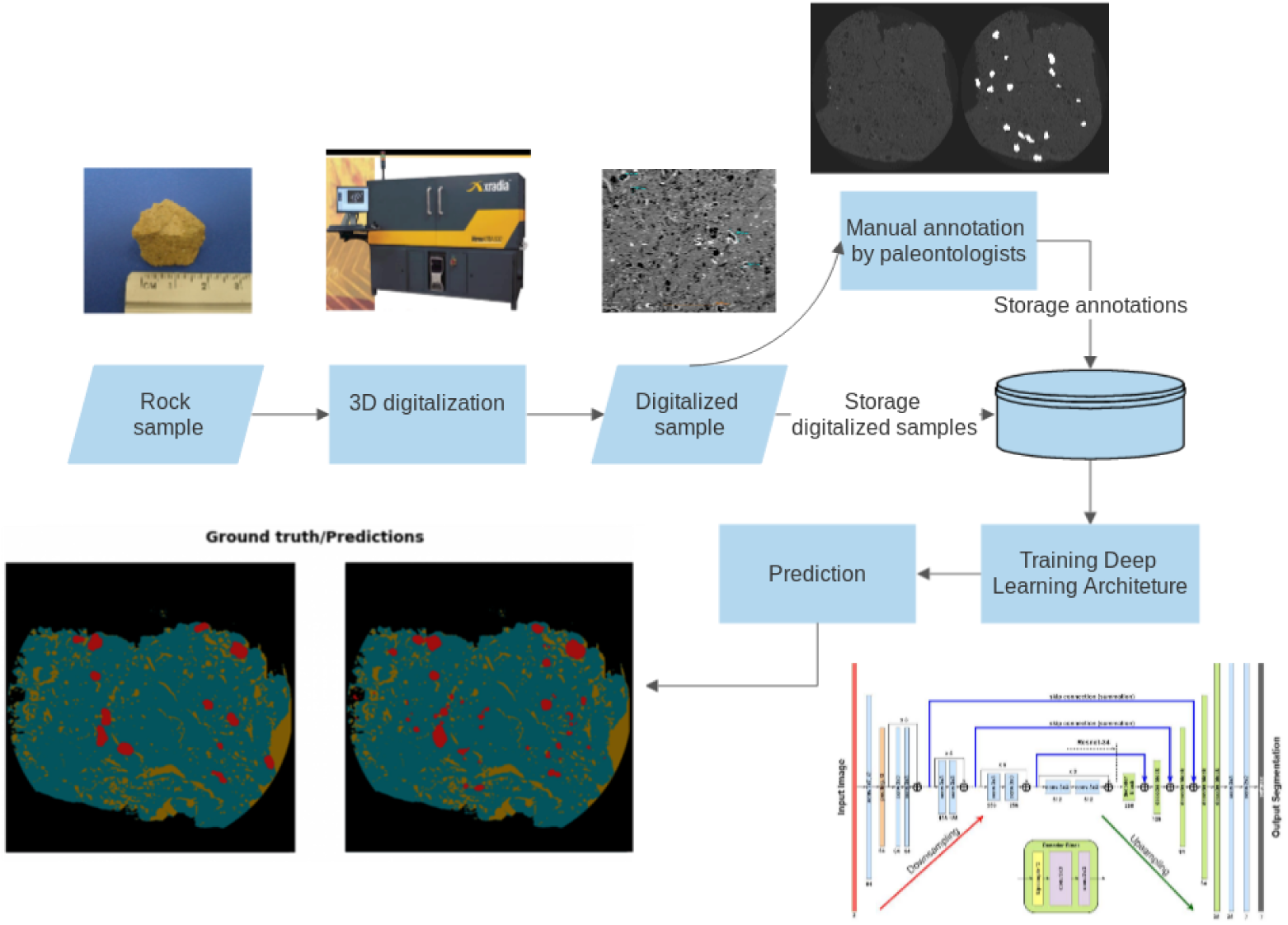
General workflow. Source: the authors and [18].

#### Non-CNN Computer Vision Methods

As part of a prospective search for CV methods for the microfossils segmentation, before we started investigating the use of CNNs, we performed a series of experiments using non-CNN, *i.e.* conventional CV methods for the segmentation of the MicroCT volume.

We analysed an extensive list of conventional CV algorithms, searching for a segmentation algorithm which, with the most appropriate input parameters, would potentially generate satisfactory results. We identified as interesting and selected the following classical segmentation algorithms: active contours [9], simple threshold and threshold with OTSU [16], all taking into account the complete tomographic volume. In order to find the best possible parameters for each segmentation algorithm, we performed a broad parameter values search running the algorithms with varied parameter sets. For the active contour algorithm, in order to find the best parameter set, we employed a genetic algorithm to search through possible input parameters. For this purpose we considered 5 input parameters: Number of steps, Sigma, Alpha, Smoothing and Theta, in a broad range of values.

The results of these conventional CV algorithms were initially analysed through *visual inspection*. For the conventional CV method that presented the best results to the visual inspection, we subsequently analysed its results also quantitatively employing the method described below.

#### CNN-based Segmentation Methods

In the 3D object identification and segmentation field, the most successfully and commonly used SS models in the last years have been the UNET and its variations. The UNET architecture was presented in [19], where the authors show its use for medical image segmentation. UNETs provide a general framework that can be parameterized with a specific *image classification* CNN model. The UNET then employs two slightly modified instances of this CNN, an *encoder* and a *decoder*, one for image recognition and another, employed in reverse mode, for the segment mask generation [1]: it uses the encoder to map raw inputs to feature representations and the decoder to take this feature representation as input, process it to make its decision and produce an output. As the UNET produces state of art semantic segmentation we choose it as our starting point.

In our work, we initially employed the UNET model associated with a ResNet34 [7], as our initial structure and added several state of art improvement methods. These methods were: nearest neighbour interpolation and pixel shuffling [20], Leaky Relu [28] for activation function and batchnorm [2] for batch normalization. This complete structure is available at the fastai ^1^ framework, which is a framework over Pytorch that contains several models, methods and state of art improvements.

#### Evaluation metrics

We evaluated each employed segmentation method comparing our results to the ground-truths generated by the micropaleontologists using the Intersection Over Union (IOU) [17] score, which quantifies similarity between finite sample sets, and is defined as the size of the intersection divided by the size of the union of the sample sets. The predicted labels were evaluated against a specialist generated ground truth.

## Results

This section presents the results we obtained with the different algorithms and CNN models we tested.

### Conventional CV algorithms

The best results under the conventional CV algorithms we obtained employing the active contours method. For this method we obtained an IOU score of 20%. The obtained active contour segmentation result is shown in figure 5.

**Figure 5.**
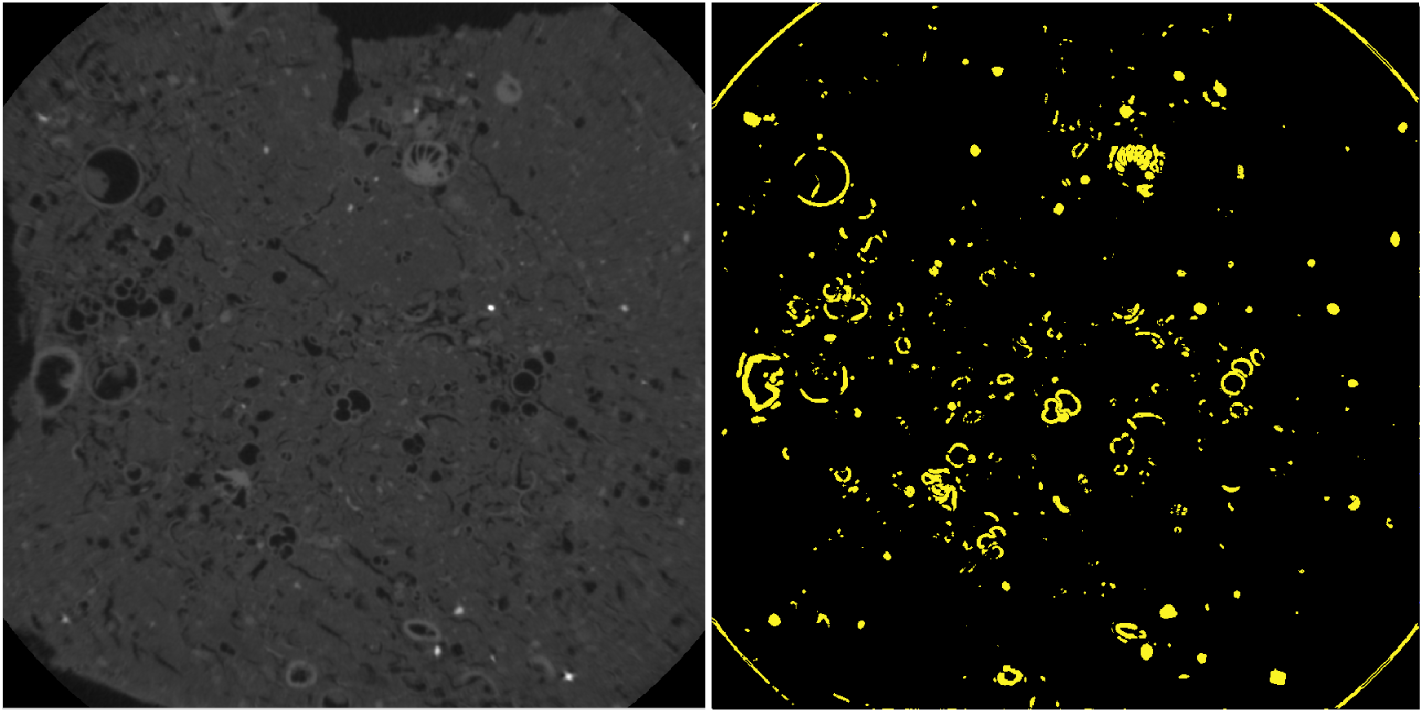
Best microfossil segmentation that we could obtain using 3D active contours (IOU = 20%). Source: the authors.

The results we obtained indicated that conventional CV methods may not be indicated for the task of microfossil segmentation in rock samples.

### CNN-based Semantic Segmentation

For our initial tests with SS CNN models, we started with the following structure: UNET associated with ResNet34 and the binary cross entropy as its loss function, a carbonatic rock sample with several microfossil specimens, scanned with the MicroCT previously described resulting in a total of 1000 slices. We employed an Intel Core i7-7700 CPU3.60GHz, 32GB memory computer and an NVIDIA GeForce GTX 1080 Ti 11GB GPU.

With this initial structure, our first experiment used only the microfossil annotation, performing a binary classification between microfossil and everything else. To improve initial results some strategies such as data augmentation and transfer learning were applied aiming to minimize the effect of having a small database. However, the obtained IOU coefficient, used for the results evaluation, stopped in 40-45%. Trying to improve the obtained result, we increased the number of classes to four, which divided the everything else class into porous space, rock and background. With this number of classes, the obtained IOU value went from 40-45 % to 75-76 % and stopped. One problem with this approach is the data balance [31], *i.e.*, the existence, in the samples, of more annotations from the rock class in comparison with the microfossils. The figure 6 shows the result obtained after marking and training for the 4 classes setup for a selected slice.

**Figure 6.**
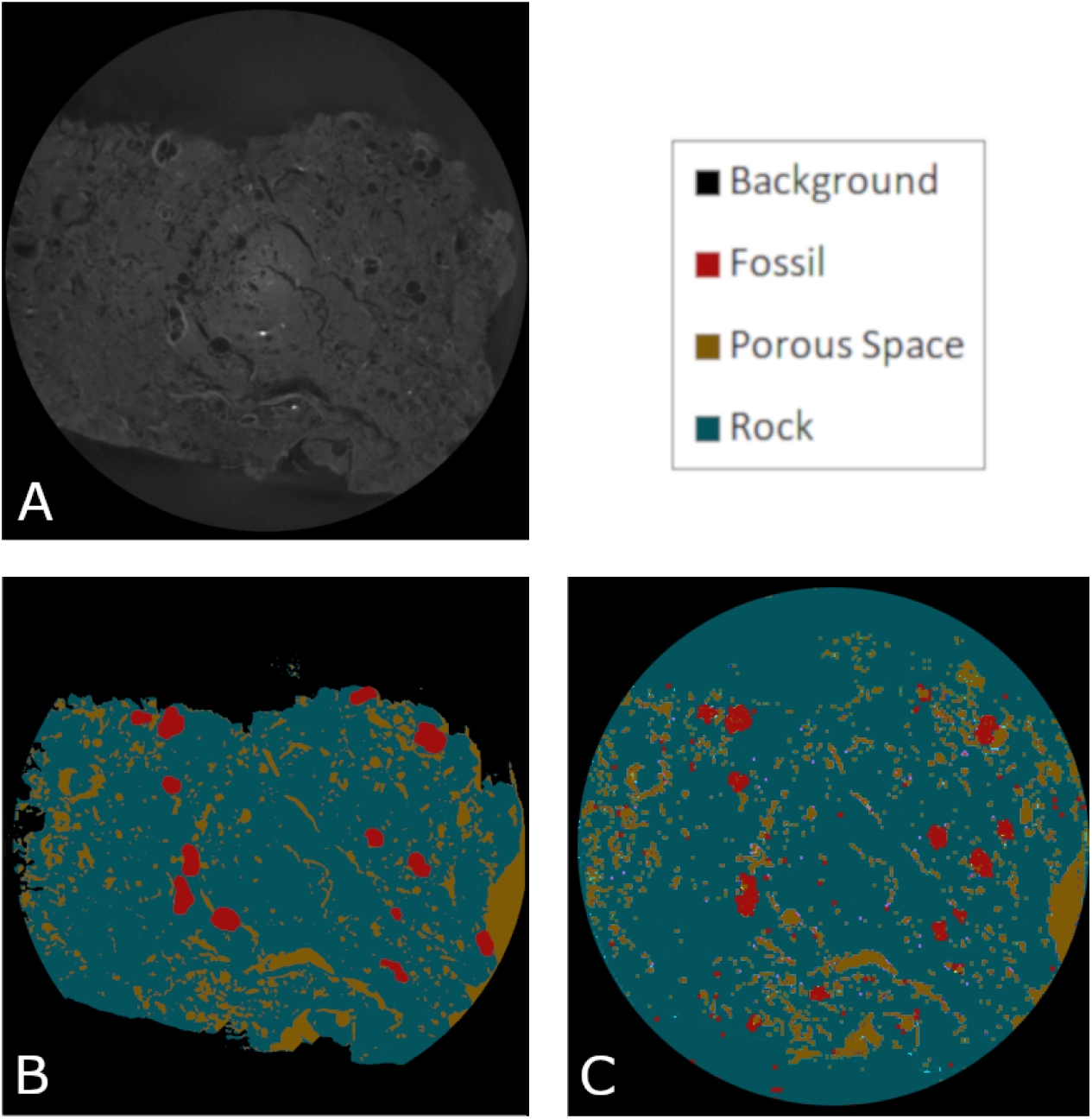
Obtained microfossil segmentation results with the 4-classes approach. (A) Original digitalised image. (B) Ground Truth manually generated by paleontologists. (C) UNET + ResNet34. Source: the authors.

Still using the 4-classes approach, we adjusted the hyper-parameters and applied a few performance-enhancing strategies [27], such as the *progressive input image resolution enhancing approach* (Jeremy Howards, informal communication during lecture at https://course.fast.ai/videos/?lesson=1) and explored data augmentation and batch size in order to obtain a 98% IOU. The microfossil GT and its resulting segmentation with this best IOU is shown in figure 7.

**Figure 7.**
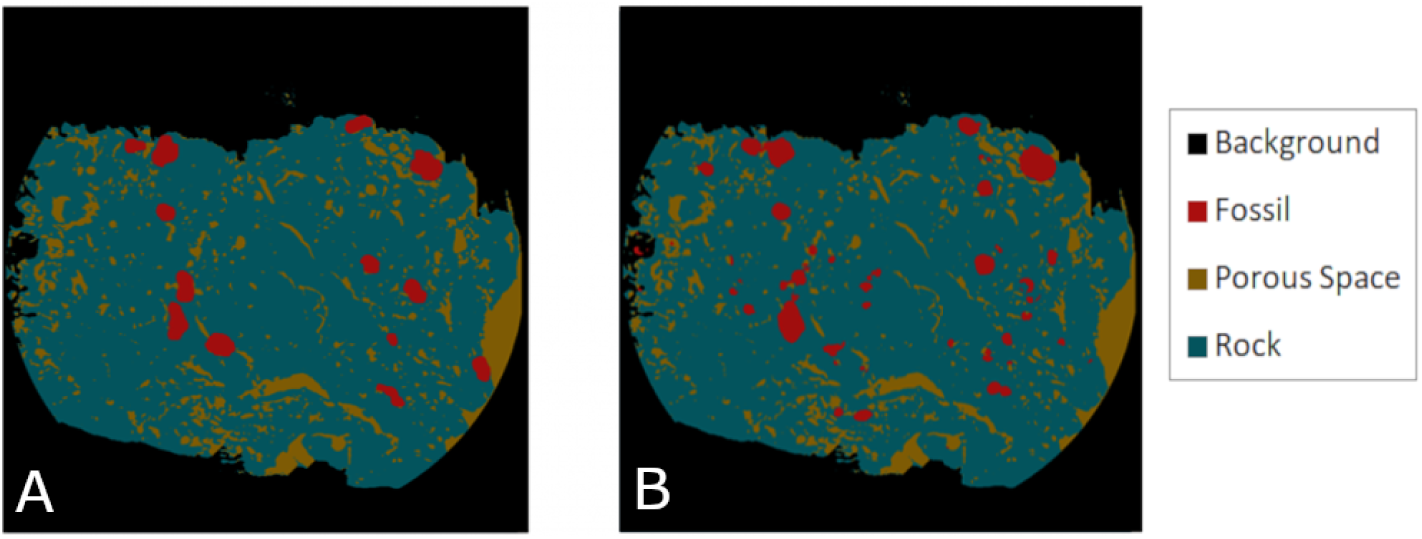
The ground truth (A) and the obtained microfossil segmentation result (B) with 4 classes, automated hyper-parameters search and additional data augmentation. Source: the authors.

Our experiments resulted in an experimental environment, where we employed the UNET as base model associated with other models in the decoder part (restnet18, ResNet34, ResNet50, ResNet101), the Cross entropy as loss function and IOU for quality assessment. Table 1 shows the IOU value for each method employed and figure 8 shows the original image, its GT and the prediction results for all architectures we tested.

**Table 1.** Segmentation performance in terms of IOU value. Each method was evaluated in a set of 1000 images from annotate microfossil data.

| Method | IOU score |
| --- | --- |
| UNET + ResNet34 | 0.76 |
| UNET + ResNet18 + hyper-parameter optimization | 0.97 |
| UNET + ResNet101 + hyper-parameter optimization | 0.97 |
| UNET + ResNet34 + hyper-parameter optimization | 0.98 |
| UNET + ResNet50 + hyper-parameter optimization | 0.98 |

**Figure 8.**
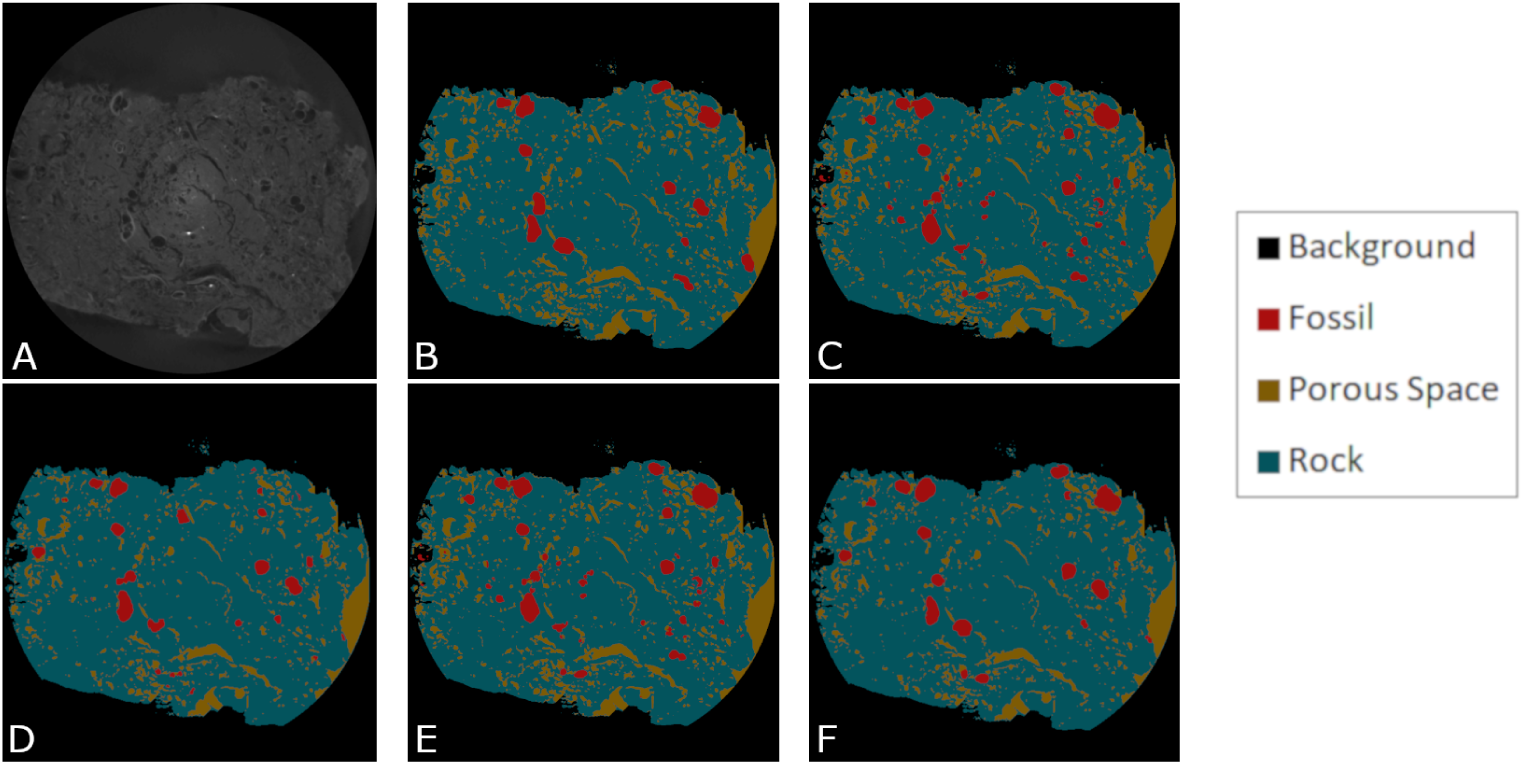
(A) Original digitalised image. (B) Ground Truth manually generated by paleontologists. (C) UNET + ResNet18 + hyper-parameter optimization. (D) UNET + ResNet101 + hyper-parameter optimization. (E) UNET + ResNet34 + hyper-parameter optimization. (F) UNET + ResNet50 + hyper-parameter optimization Source: the authors.

After segmenting we took the predicted mask for resnet34 and applied to the original image. The result of this process is the easy identification of several microfossils. Figure 9 shows the mask overlap result, the identification of one microfossil specimen (highlighted with the red rectangle) followed by its magnified version and the correlation of this the magnified version with other two versions of the same specimen (physically isolated and digitized with the Versa XRM-500 MicroCT and the Zeiss Discovery V20 stereoscope).

**Figure 9.**
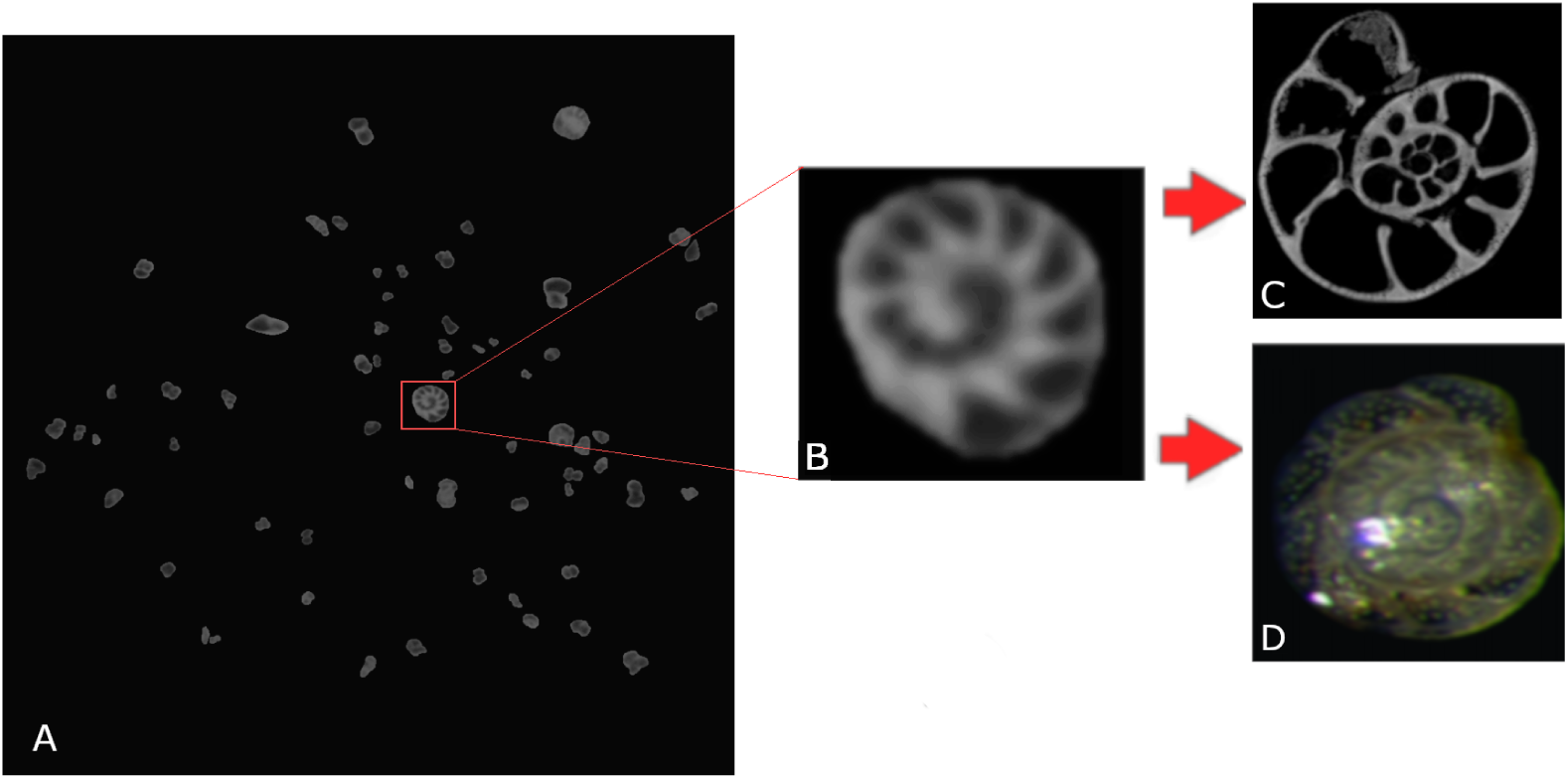
Result of applying the obtained segmentation mask over the digitized image. (A) contrast-enhanced 2D section image masked from the digitized MicroCT volume with one specific microfossil highlighted in red. (B) Highlighted microfossil extracted and magnified for visualisation. (C) Physically isolated microfossil digitized with The Versa XRM-500 MicroCT. (D) Cibicidoides multidimensional acquisition with the Zeiss Discovery V20 stereoscope. Source: the authors.

## Discussion and Conclusions

In this paper we present a new nondestructive processing pipeline for the identification of microfossils in carbonatic rocks that allows for the fully automated segmentation of these fossils without the need of previous physical separation. Furthermore, we developed and validated the CV methods for this identification and segmentation. The validation was quantitatively and automatically performed against a ground truth manually generated by expert micropaleontologists.

An extremely relevant aspect of the developed pipeline for the field of paleontology, more specifically micropaleontology, resides in the nondestructive character of the method. In the micropaleontological study process an essential step is the samples preparation, aiming to separate the microfossils from the other rock and/or sediment constituents. In the traditional laboratory process, the samples are physically disaggregated (ground or milled) and subsequently performed chemical disaggregation, with addition of reagents (e.g., hydrogen peroxide and acetic acid). Both physical or chemical disaggregation can alter or even destroy microfossils characteristics. In this premise, the imaging method is crucial for the morphological characteristics visualization as reliable as possible, allowing the individuals taxonomic recognition [8].

Another relevant factor that makes this method interesting is that it allows the microorganisms’ preservation analysis throughout geological time, as well as aspects of fossilization, preservation and even position in which the microfossils are deposited (preserved) in the rocks. It should be emphasized that studies with the taphonomic approach are fundamental for paleoenvironmental conditions and/or diagenetic alteration processes reconstitution over geological time. Also, the use of this tool is strongly indicated in cases where it is extremely difficult to recover microfossils along specific sections and/or intervals where the material (rock) is very compact and even when it presents incipient diagenetic alteration. The microfossils identification is strategic for the exploration of petroleum due to the use in biostratigraphy, which refers to the use of microfossils from different groups to perform the temporal characterization of sedimentary rock strata, fundamental for the petroleum industry.

A few observations can be performed from the obtained results: (i) the importance of employing appropriate hyper-parameters such as learning rate, weight decay, momentum and batch size: with that hyper-parameters optimization we obtained an improvement of 20 %. (ii) a network architecture grown does not imply in better results. It is possible to observe that the ResNet34 shows the same results that the ResNet50 and a better result when compared with a ResNet101. However, here we have a hardware limitation: both, ResNet50 and ResNet101, couldn’t run with the full image resolution on the 11 GB GeForce, even with a batch size of 1. Even so, the ResNet34 requires less execution time and hardware. (iii) Analyzing the obtained result images and comparing the visually against their Ground Truth (Figure 8), we still notice some small errors, however, we understand that this can be be mitigated by adding more training samples, together with GTs from experts, to the training set when applying this pre-trained network to other, new, samples. Also, there are always new state of art improvements that could by tried aiming to reduce even more this small errors. Figure 9 shows the isolated microfossil digitalized and its correlated identification into the sample.

We understand that this process of microfossil identification without the need of physically isolate the microfossil has the potential to allow the paleontologist to analyze specific aspects of a sample such as the microfossils deposition. This is important for some applications in the oil and gas industry. It also has the potential to improve the paleontologist’s work, because instead of losing time to physically isolate the microfossil he receives the microfossil already identified and can perform other analysis such as class identification and orientation.

### Threats to validity

We employed a dataset that, even if it consisted of a very large quantity of images and presented a wide variety of microfossils, was gained from a sample obtained from a singular drill probe. On the other side, the samples digitisation and annotation afford a set of requirements such as: having a MicroCT working and available; the cost of the MicroCT digitisation process; a storage to keep the amount of generated data; and a paleontologist group to analyze and annotate each digitised sample. As the workflow we suggest in this paper is new, it was not in place on any of the partners that participated in this work and to obtain more scanned and annotated samples was not possible at this point of our research.

This could jeopardize the generalizability of this work, as we have not enough data to claim that our approach will be successfully applicable to any carbonatic rock sample. On the other side, our identification and segmentation results were extremely successful and we understand that they are promising. From the authors’ knowledge there is not any other publicly available carbonatic rock probe dataset, with or without specialist-annotated microfossils.

In this context, we understand our work as pioneering and pointing to a promising direction of research that can potentialize both, micropaleontological research and associated economical activities, such as oil prospection. Our publicly available fully annotated MicroCT database has also the potential to support research activities to be performed by other groups.

### Conclusions

Summarizing, this work presents:

- the first fully annotated MicroCT-acquired publicly available microfossils dataset;
- a baseline for microfossil segmentation and the comparison with deep learning-based semantic segmentation and other segmentation architectures;
- a methodology for microfossil studies through MicroCT-acquired digital models;
- a tool for cases where it is extremely difficult to recover microfossils along specific sections.

With the improvement in the available hardware some future works aim to reduce even more the obtained errors by increasing the batch size and image resolution and employ more state of art deep learning improvements.

## Acknowledgments

This study was financed in part by the Coordenação de Aperfeiçoamento de Pessoal de Nível Superior - Brasil (CAPES) - Finance Code 001 and by PETROBRAS through the research project number 902. There are no conflicts of interest.

## Footnotes

1 https://www.fast.ai/

